# Smartphone deprivation alters cortical sensorimotor processing of the hand

**DOI:** 10.1101/2021.03.04.433898

**Authors:** Arko Ghosh

## Abstract

Brain areas representing the body can change in response to behavioral alterations. This idea is firmly established for the adult cortex in response to extraordinary alterations such as traumatic amputation or casting of the limb. Here we address how adult cortical sensorimotor processing alters in response to a subtle perturbation in the form of smartphone deprivation lasting for ~7 days. We quantified the sensorimotor processes associated with the fingertips before and after the deprivation in right-handed smartphone users. The measurements were contrasted with those of a control group with unperturbed smartphone behavior. First, smartphone tapping speed in daily life became slower after the deprivation. Second, according to reaction time tests conducted in the laboratory the asymmetrically superior performance of the right vs. left thumb was eroded by the deprivation. Third, according to EEG measurements at physical rest, tactile stimulation at the right thumb tip resulted in smaller signal amplitudes after the deprivation. Moreover, the EEG measurements during smartphone use revealed larger signal amplitudes for tactile stimulation at the right little fingertip after the deprivation. We show that cortical plasticity can occur by merely disengaging from a common day-to-day behavior. We suggest that in daily life the adult brain continuously and selectively updates its sensorimotor processing according to recent experience.

## Introduction

Alterations in body use can drive adult brain plasticity of the corresponding sensory and motor cortices. For instance, after traumatic amputation, there is a considerable reorganization of the cortical areas associated with the missing and compensating limbs ^1–4^. Such plasticity is not limited to severe injury but can also occur due to immobilization of a broken arm^5^. Moreover, experimental constraints of the limb in both human and non-human primates cause substantial changes to cortical sensorimotor processing ^6–10^. A consistent finding across these diverse explorations is that the cortical processing associated with the disused limb is diminished. In daily life, a reduction in limb use may occur due to shifting behavioral demands. In modern behavior, shifts in thumb use may be driven by the fluctuations in smartphone usage and this ubiquitous behavior can be seamlessly captured in the background by using server logs, or sensors embedded in the smartphone ^11–13^. Despite the seamless behavioral monitoring possibilities we know little about how the brain adapts in response to a momentary shift in smartphone usage.

A distinct line of research on use-dependent plasticity is focused on increased use rather than disuse of the limb^14^. For instance, according to observational studies in string instrument players the amplitude of cortical sensorimotor signals evoked from the little finger is correlated to musical practice^15^. This idea of use-dependent plasticity is further supported by intervention studies where users gain new skills ^14^. For instance, learning to juggle introduces structural alterations in the cortical sensorimotor areas in contrast to an untrained controlled group^16^. Such plastic processes may not be confined to exception skills given that smartphone users also show enhanced cortical processing from the thumb compared to people without smartphones^17^. In particular, the middle to late latency somatosensory signals (60 to 150 ms) evoked from the thumb – but not the signals evoked from the fingers less used on the smartphone – are correlated with smartphone skills and recent fluctuations in smartphone usage^17,18^. The correlates are not confined to neurophysiological measures as faster smartphone interactions in daily life have been found to lower the trial-to-trial motor variability on a tactile reaction time task administered in laboratory ^18^.

Although the acquisition of exceptional skills and their corresponding sensorimotor cortical correlates are well documented, the neuronal consequences of disusing a skill are not clear. More specifically, it is not clear how the brain adapts in response to the disuse of a skill that has been “crystallized” through repeated usage in daily behaviour^19^. According to a meta-analysis of behavioral data contrasting natural skills such as typing versus artificial skills acquired for laboratory testing, natural skills are less susceptible to decay than their artificial counterparts in response to disuse^20,21^.

To empirically address if and how disuse of a common behavior impacts cortical sensorimotor processing we deprived willing volunteers of their smartphones for ~7 days. We leveraged the smartphone touchscreen logs to quantify the extent to which subjects followed the deprivation instruction. The background App also allowed us to assess the impact of the deprivation on the sensorimotor touchscreen skill expressed in daily life. In the laboratory, behavioral testing and EEG measurements showed that cortical sensorimotor processing from the hand was significantly altered after the deprivation.

## Results

### Impact of smartphone deprivation on touchscreen interactions

According to self-reports, participants preferentially used their right hand on the smartphone touchscreen (**Fig. 1**). This was consistent with the sampling strategy as only self-reported right-handers were recruited. The thumb was the preferred choice in virtually all of the participants while the little finger was the least preferred (**Fig. 1**).

**Figure 1.**
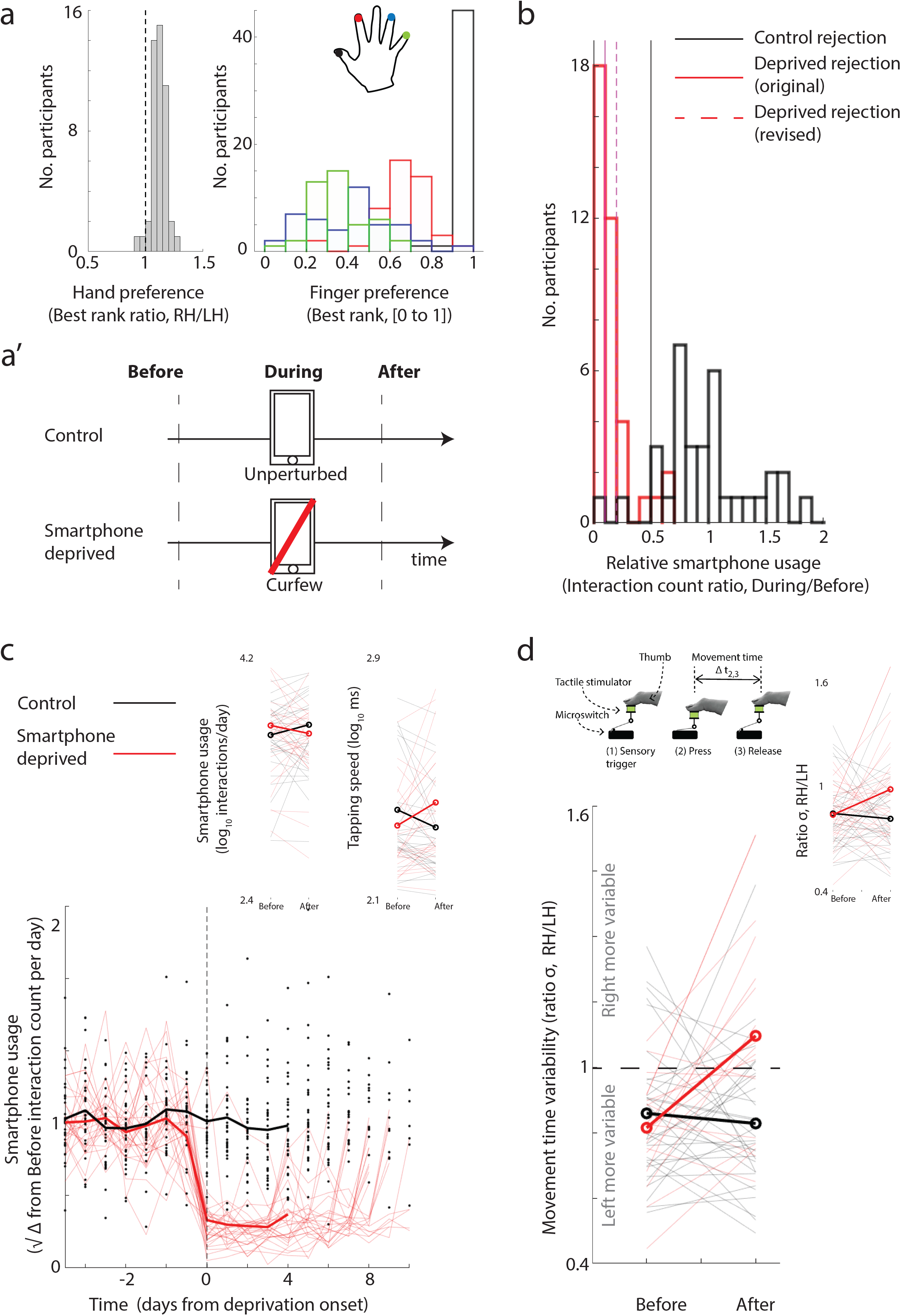
Smartphone deprivation and its behavioral impact. **(a)** The sampled population predominantly reported that the right hand and the thumb were the most used on the smartphone touchscreen. **(a’)** We deprived individuals of their smartphones and performed EEG and behavioral measurements before and after the deprivation. A control group received no instructions to alter smartphone behavior. **(b)** The ratio of typical (median) smartphone usage between the two measurement times and the usage before the first measurement. Note, the instructed targeted usage for the deprived group was 0.1 (10% of their typical usage based on the previous 21 days) but for the analysis presented here users reaching up to 0.2 were included. Users in the control group who dropped below 0.5 were removed from further analysis. **(c)** The deprivation instruction had a sustained impact on smartphone usage through the corresponding period. The population (mean) usage is plotted using the solid lines for the two groups and thin lines show individual participants in the deprived group. Note, how the participants return to pre-deprivation levels at the end of their respective deprivation period. The first insert shows the number of touchscreen interactions generated 2 days before the deprivation and 2 days after the deprivation. The second insert shows tapping speed based on the smartphone interactions in the same period. **(d)** The motor variability measured on a reaction time task was altered by the smartphone deprivation. The same analysis including the revised deprivation target (insert).

Of the 75 participants considered here, randomly chosen 39 participants were instructed to reduce their smartphone usage to a maximum of 10% of their normal use as determined based on their ~21-day interaction history (**Fig. 1**). Given the rare nature of this approach and the heavy reliance on smartphones in daily life, we systematically quantified the compliance to this deprivation instruction. Eighteen of the 39 individuals were successful in adhering to this instruction while another 12 managed to reduce up to 20% of their normal use. The 9 remaining grossly incompliant participants were eliminated from further analysis. The 36 individuals in the control group were required to maintain their activity down to a minimum of 50% of their normal use in between the two measurements – although they were not explicitly instructed to do so. Three of them fell below this threshold and were eliminated from further analysis. The uneven number of eliminations between the two groups suggests an effective sampling bias towards individuals who were more accepting of not using their smartphones in the deprivation group.

The deprived group showed a substantial drop in usage from a pre-deprivation mean of 4738 ± 1913 SD interactions per day to 403 ± 242 SD interactions per day during the deprivation. The control group remained relatively unchanged in the same period, from a mean of 4114 ± 2403 SD interactions per day to 3935 ± 2376 SD (*F* (1,61) = 112.72, *p* = 1.7 × 10^−15^, group × time interaction, repeated-measures ANOVA, **Fig. 1**).

Of the 63 participants, the background app remained on in all but 7 participants even after the deprivation period allowing us to assess the behavioral impact of the deprivation on smartphone use in daily life. When we compared the smartphone usage 48 hours before to the deprivation with that of the usage 48 hours after the deprivation and we found no significant change introduced by the deprivation (*F*(1,54) = 2.38, p = 0.13, group × time interaction, **Fig. 1**). However, the tapping speed showed a substantial decline in the same period (*F*(1,53) = 12.22, *p* = 0.0009, group × time interaction, **Fig. 1**, note in one subject had the insufficient number of interactions for the speed estimation).

Sleep is impacted after severe behavioral alterations such as in amputations or constrained limbs ^22^. To address if the smartphone deprivation had a broad behavioral footprint, we measured sleep using wearables. Actigraphy based sleep measures revealed that the deprivation did not impact sleep (*F* (1,56) = 0.17, *p* = 0.68, group × time interaction).

### Impact of smartphone deprivation on motor variability of the thumb

Due to the strong correlational evidence linking smartphone use and motor variability at the thumb, we addressed how the deprivation impacted the trial-to-trial motor time variability (σ of the ex-gaussian fit) during a tactile reaction time task^18^. We used the ratio of the variability obtained from the right thumb versus that of the left thumb as an index of variability ‘normalized’ to the control hand that is seldom used on the smartphone. We analyzed the impact of deprivation on this measure for the participants who met the original criteria of deprivation (10% of typical use) and for the larger group of participants who met the revised criteria (20% of typical use, **Fig. 1**). In both the control and deprived groups, the measure was pre-dominantly below 1, reflecting a pre-existing higher motor variance of the left thumb than of the right (42 of 51 participants who met the original criteria, and 50 of 63 who met the revised criteria). The index of variability increased after smartphone deprivation and this was evident in both the smaller group who met the original deprivation criteria (*F* (1, 48) = 13.22, *p* = 0.0007, group × time interaction, **Fig. 1**) and the larger group who met the revised criteria (*F* (1, 60) = 5.17, *p* = 0.027, group × time interaction, **Fig. 1**).

Although the left thumb offered a point of reference in the above analysis, the impact of deprivation may be detectable in the unreferenced data. Essentially, the deprivation-induced increase of the variability index may be attributed to reduced variability at the left thumb (denominator of the variability index) or increased variability at the right thumb (the numerator of the variability index). Alternatively, in the case of a mechanism that depends on an interaction between the two sides (to maintain the level of left-right asymmetry), left and right separated measures would be less likely to reveal significant alterations after the deprivation. Therefore, we conducted follow-up statistical tests on the left and right thumb variability measures separately. There was no consistent impact of the deprivation on these measures (left thumb, original criteria, *F* (1, 48) = 3.078, *p* = 0.086, revised criteria, *F* (1, 60) = 2.39, *p* = 0.127; right thumb, original criteria, *F* (1, 48) = 2.21, *p* = 0.14, revised criteria, *F* (1, 60) = 0.34, *p* = 0.56, group × time interaction).

### Somatosensory evoked potentials at physical rest

Tactile stimulation of the right thumb at physical rest resulted in early signals over the contralateral (left) cortex. We detected distinct signal peaks at 20 ms, 50 ms, 85 ms, and 200 ms measured over a midline frontal (Θ = 45, Ф = 90) electrode (**Fig. 2**). Smartphone deprivation suppressed the amplitude of the mid latency components ranging from 75 ms to 135 ms. Statistically significant clusters showing a group × time interaction were found over the frontal and central electrodes (**Fig. 2**, repeated measures ANOVA performed at each electrode and time points between −200 to 400 ms from the stimulation followed by multiple comparison correction, N = 49, signal rejections based on amplitude baselines). The same pattern of results but with enhanced *F* values was found when the pre-deprivation smartphone usage, the tapping speed, the inter-measurement gap, and the gender were taken into consideration as covariates in a linear model (**Fig. 2**, ANCOVA based on gain scores derived from pre & post measurements. The test was performed at each electrode and time point between −200 and 400 ms from the stimulation, N = 53, signal rejections based on gain score reference at baseline). The deprivation did not impact the cortical responses to tactile stimulation of the right little finger (**Fig. 2,** ANOVA N = 50, ANVOCA N = 52), the left thumb, or the left little finger (**Suppl. Fig. 1,** ANOVA N = 45 & 47, ANCOVA N = 49 & 52). A nearly identical pattern of results was obtained upon the inclusion of only those participants who were right thumbed on the smartphone according to self-reports (**Suppl. Fig. 2**, ANOVA N = 38).

**Figure 2.**
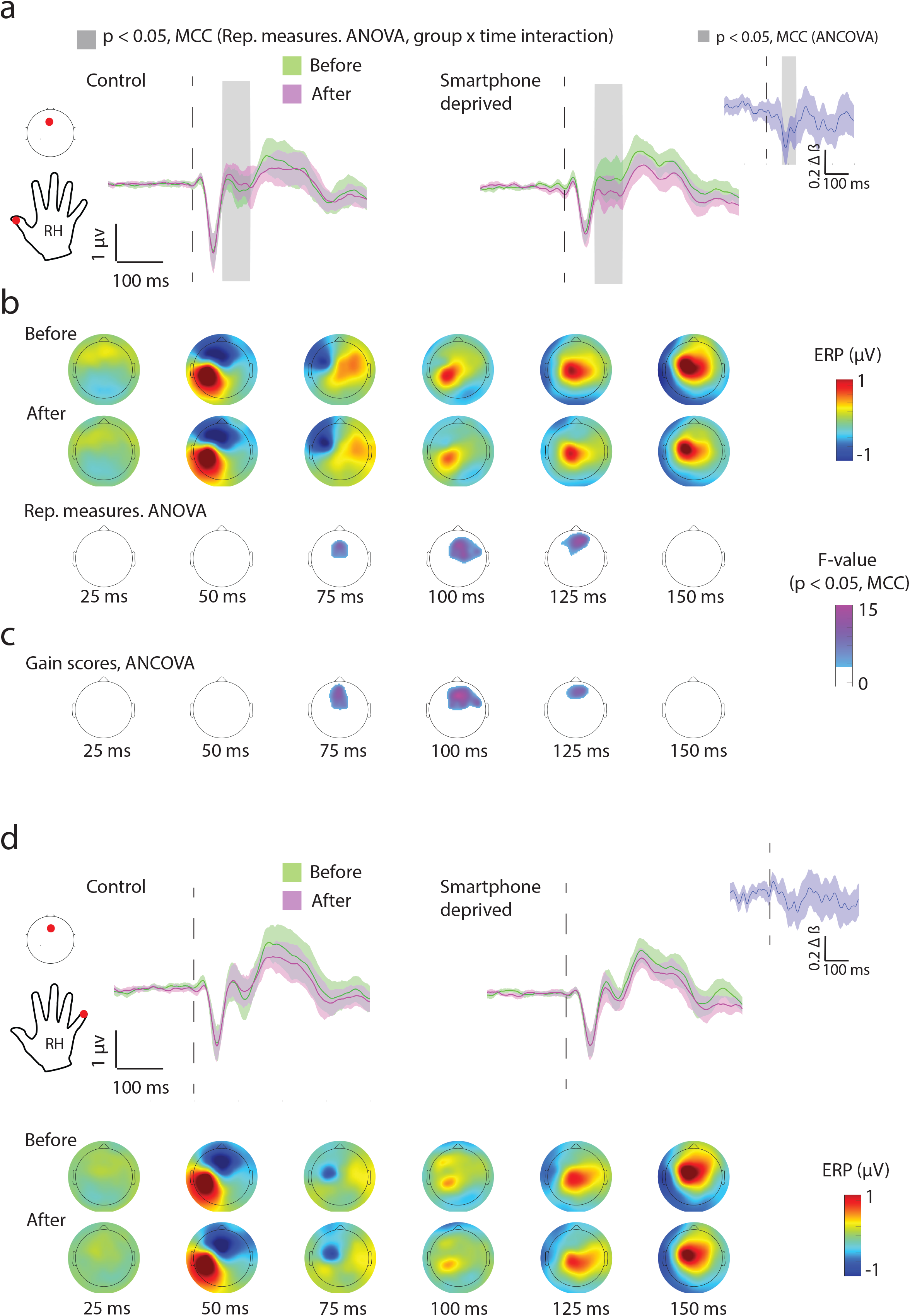
Event-related potentials corresponding to tactile stimulation of the right hand at physical rest. **(a)** Tactile stimulation at the thumb tip results in smaller amplitude EEG signals after smartphone deprivation. Mean (trimmed) and the corresponding 95% confidence intervals are shown from a midline frontal electrode (red dot) with the period of significant group × time interaction clusters (according to repeated measures ANOVA corrected for multiple comparison correction, abbreviated as MCC) shaded in grey. A parallel set of analyses was performed using ANCOVA based on pre to post measurement gain scores. The inserted plot shows the difference in the ANCOVA model’s *β* values for the control group vs. the deprivation group. **(b)** Scalp plots show mean signals from the smartphone deprivation group and their corresponding interaction *F*-values from the ANOVA. **(c)** Corresponding *F*-values of the ANCOVA model for the differences between the inter-measurement session gain scores between the control group vs. the deprivation group. **(d)** Tactile stimulation at the little finger did not reveal any significant alterations after smartphone deprivation. See Supplementary Figure 1 for results from the left hand. See Supplementary Figure 2 for results from a subset of volunteers with the non-right thumbed smartphone users excluded.

### Somatosensory evoked potentials during smartphone use

Before addressing if the sensory alterations were detectable in a distinct condition – where the participants were on their smartphones as opposed to physical rest – we first characterized how the sensory signals differed between these two conditions. Towards this, we compared the EEG signals evoked by the tactile stimulation in the passive condition with during smartphone use. For both the thumb and little finger we found suppressed signal amplitudes in the smartphone use condition (**Suppl. Fig. 3,** Thumb N = 47 & Little finger N = 50). The suppression was significant in the early to the long latency signal components. Next, we addressed whether engaging on the smartphone distinctly impacted the sensory processing of the thumb versus the little finger. The engagement-induced depression was more pronounced for the thumb and statistically significant in the mid (40 ms to 120 ms) and long (170 ms to 280 ms) latency signal components (**Suppl. Fig. 3**, stimulus location × condition interaction, repeated measures ANOVA). In summary, tactile sensory processing during smartphone use was distinct in comparison to the passive condition and the suppression in the smartphone use condition was sensitive to the location of the tactile input.

Next, we addressed how the deprivation impacted tactile processing during smartphone use. The EEG signal amplitudes corresponding to the right thumb remained unchanged (ANOVA N = 52, ANCOVA N = 56). Interestingly, the mid latency signal amplitudes (50 ms to 125 ms) corresponding to the right little finger were enhanced after the deprivation (**Fig. 3**). According to a repeated-measures ANOVA, significant group × time interaction clusters were detected at the central and frontal electrodes (**Fig. 3,** ANOVA N = 58, ANCOVA N = 58). A similar pattern emerged using the ANCOVA gain scores analysis. Besides, the ANCOVA revealed clusters at the temporal electrodes contralateral to the stimulation (**Fig. 3**).

**Figure 3.**
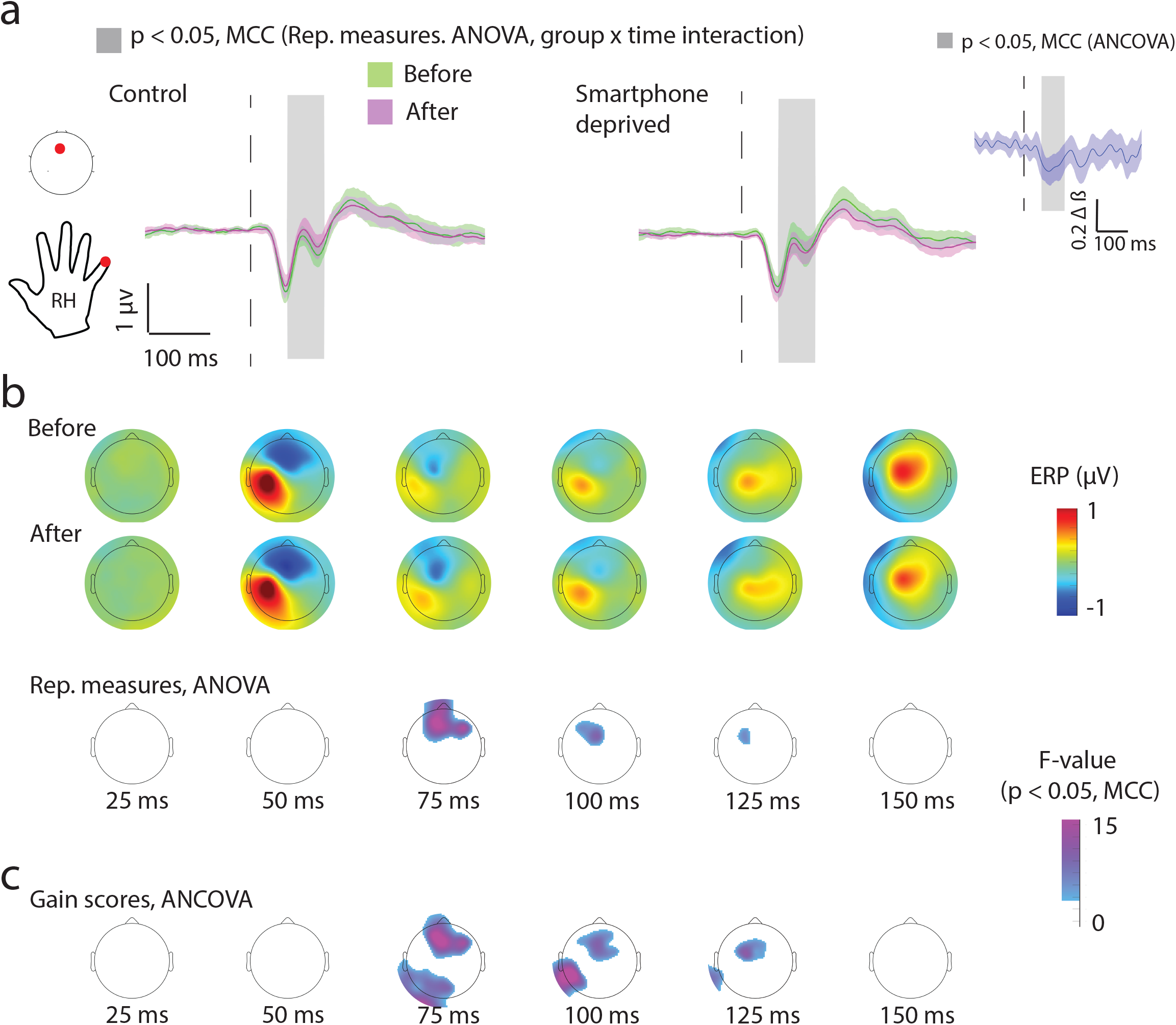
Event-related potentials corresponding to the right little finger evaluated during smartphone use. In contrast to the physical rest condition, in this condition, the participants were actively using their smartphones during the presentation of the tactile stimulation. The EEG measurements revealed higher amplitude signals upon stimulation at the little fingertip after the deprivation. The figure attributes are the same as in Figure 2. **(a)** The signal amplitudes at the midline frontal electrode. Insert, the difference in *β* values derived from the ANCOVA model. **(b)** Scalp plots of mean signal amplitudes. The corresponding *F* values for the repeated measures ANOVA interaction. **(c)** The *F* values obtained by using ANCOVA (based on the gain scores). See Supplementary Figure 3 for a comparison between tactile event-related potentials at physical rest versus during smartphone use.

### Repetition suppression between measurement sessions

A common observation in neuronal measurements is that the neuronal signals are reduced in amplitude when the stimuli are repeated within the same experimental session^23^. In our study, participants experienced the same bout of tactile stimuli in two sessions separated by several days. We addressed whether the second session showed suppressed inputs compared to the first using the ANOVA models as presented above but by focusing on the main effect of time. Under the condition where the participants were at physical rest, all of the stimulated fingertips showed significant clusters for the main effect of time (**Suppl. Fig. 4**). Interestingly, the significant clusters were found only for mid to long latency EEG signal components (> 110 ms). Moreover, when stimulations occurred during smartphone use, we found no significant cluster reflecting the main effect of time. Notably, in none of the ANOVA models did we ever find a significant cluster reflecting the main effect of the group.

## Discussion

In the present study, a period of partial smartphone deprivation altered sensorimotor cortical functions associated with the hand. The deprivation slowed the smartphone tapping speed in daily life. In the laboratory, we established that the deprivation eroded the asymmetry in movement timing variability between the two thumbs. Moreover, the EEG signal amplitude corresponding to the tactile stimulation at the right thumb tip was depressed after the deprivation while the signals corresponding to the right little fingertip were enhanced. The overall pattern of changes induced by smartphone deprivation is consistent with the ideas of use-dependent sensorimotor modifications established by the study of limb injury or limb immobilization. Still, our findings suggest a distinct form of plasticity that is triggered by the disuse of a specific behavior as opposed to the general loss of limb functions.

Under conventional frameworks of motor learning highly practiced behaviors are considered as “crystalized” and may be reliant on neuronal representations stabilized by practicing ^14,24,25^. However, our data showing that smartphone tapping speed evaluated in the first 48 hours after the deprivation is slower than the pre-deprivation speed suggests that there are limitations to this stabilization. This pattern of results makes it likely that the smartphone tapping skills are actively maintained by day-to-day touchscreen interactions.

In general, the use of the dominant hand may developmentally shape the corresponding cortical circuits to generate less variable movements than the non-dominant hand ^26–28^. Interestingly, performance asymmetry is diminished in right-handed musicians and this is attributed to developmental adaptations during the acquisition of musical skills requiring bi-manual coordination^29^. Our findings of reduced asymmetry in motor variability after smartphone deprivation support the role of daily behavior in shaping the corresponding circuits into adulthood. We could not attribute this reduced asymmetry to increased variability of the right thumb or reduced variability of the left thumb suggesting that the underlying cortical processes driving the asymmetry are shaped by inter-hemispheric interactions. Alternatively, we must consider these findings under the assumption that the circuits underlying the left thumb are unaltered by the deprivation and that they serve to normalize the right thumb measures to general motor variability of the hands. According to this consideration, our findings indicate that the deprivation increases variability in cortical circuits associated with the right thumb^18^.

The EEG measurements in response to tactile stimulation aided at two levels in our understanding of the deprivation-induced sensorimotor cortical plasticity. First, they helped isolate which body parts were prominently impacted by the deprivation; and second, they helped address which stages of tactile information processing were susceptible to the perturbation. In the passive condition – in which the participants were at physical rest – of the four stimulated locations (thumb and little finger of both hands) only the right thumb altered with the deprivation and the corresponding EEG signal amplitudes were depressed. This change is in keeping with correlational evidence that the right thumb is uniquely sensitive to recent smartphone experiences ^17,18^. Moreover, the changes were limited to the mid-and late-latency somatosensory signals. As these signals likely originate from the secondary somatosensory cortex, our findings suggest this higher sensory area is fine-tuned by subtle alterations in daily life^30^. Notably, conventional studies on somatosensory plasticity – say after loss of limb or finger function – largely reveal alterations in the primary somatosensory cortex with some exceptions extending to the secondary somatosensory cortex ^31^.

There was evidence for another form of experience-dependent plasticity in parallel to the plasticity attributed to smartphone deprivation. Essentially, tactile processing in the second measurement session was suppressed when the processing was evaluated at physical rest. This is akin to the previously documented repetition suppression between experimental sessions separated by days and observed in the visual system^23,32,33^. Interestingly, only mid and long-latency signals were suppressed and thus the suppression was encoded at a distinct processing stage than the smartphone deprivation-induced alterations. This raises the possibility that the experience-dependent plasticity corresponding to passive sensory processing engages distinct neuronal processing stages than the plasticity induced by momentary behavioral shifts.

We gathered evidence for the modulation of cortical tactile somatosensory processing when participants were on the smartphone. In general, somatosensory processing is suppressed when the brain performs movements in comparison to at physical rest – as if to deal with the barrage of information resulting from the movement itself ^34,35^. Consistent with this idea, EEG signal amplitudes evoked from the right thumb tip were smaller when engaged on the smartphone in comparison to when at rest. Interestingly, the amplitudes were also suppressed for the little finger – although this finger only played a supporting role to stabilize the smartphone grip. Nevertheless, this suppression was less pronounced than for the thumb tip. These results demonstrate a fine-grained suppression of sensory information that is dependent on the location of the corresponding tactile sensors.

We addressed the impact of smartphone deprivation on tactile processing during smartphone use. Interestingly, the EEG signal amplitudes from the right thumb in this condition were not altered by the deprivation. This may be an outcome of the strong suppression of signals resulting in a masking of the impact of the deprivation. The masking may have also obliterated the repetition suppression effect in the more active condition as that only occurred at physical rest. However, the deprivation did alter the EEG signals from the little finger during smartphone use. The mid-to-late- latency EEG signal amplitudes were larger after the deprivation. This is akin to the findings in the rat model of somatosensory plasticity where whiskers are selectively trimmed and the processing from the spared whiskers is potentiated ^36^.

Our study suggests that adaptations in the sensorimotor system are pervasive. Firstly, long periods of spontaneous smartphone deprivation in daily living may be rare but do occur according to the heavy-tailed inter-event distributions ^37^. Secondly, in previous empirical paradigms to study disuse induced plasticity, limb movements were suppressed but our findings show that mere withdrawal of behavior is sufficient to alter the corresponding sensorimotor circuits. Notably, conditions like amputation or constrained limbs may have a behavioral footprint that goes well beyond limb use to disrupt broad behavioral indicators such as sleep^22^. In this study, we found no evidence for smartphone deprivation impacting sleep. In sum, our findings raise the possibility that the mechanisms of plasticity are driven by specific behavioral demands in addition to the general disuse of a body part.

In conclusion, this study presents an important step towards understanding brain adaptations in daily life. Modern human behavior is dominated by smartphone touchscreen interactions and our findings underscore that this behavior is well embedded in the cortical circuits associated with the hand. This empirical approach of addressing brain plasticity is an important parallel to the conventional approaches focused on extraordinary skills or general dysfunction of the limb.

## Acknowledgments

The author would like to thank the students of the applied cognitive psychology MSc program at Leiden University who aided in the data collection (D. van Berkel, J. Borger, M. Dekker, E. van Duijn, K. van Eijgen, M. van Eijk, N. Grootscholten, K. Nieuwenhuizen, A. Tran, T. van Tuijl and M. van Weerwijk). The author deeply appreciates the help obtained from Sander Nieuwenhuis in editing this manuscript. Arthur Zhang and Irene Taboada helped with smartphone usage survey digitisation and analysis. We thank Reto Huber for providing the wearables during the study period.

---

**Supplementary Figure 1.**
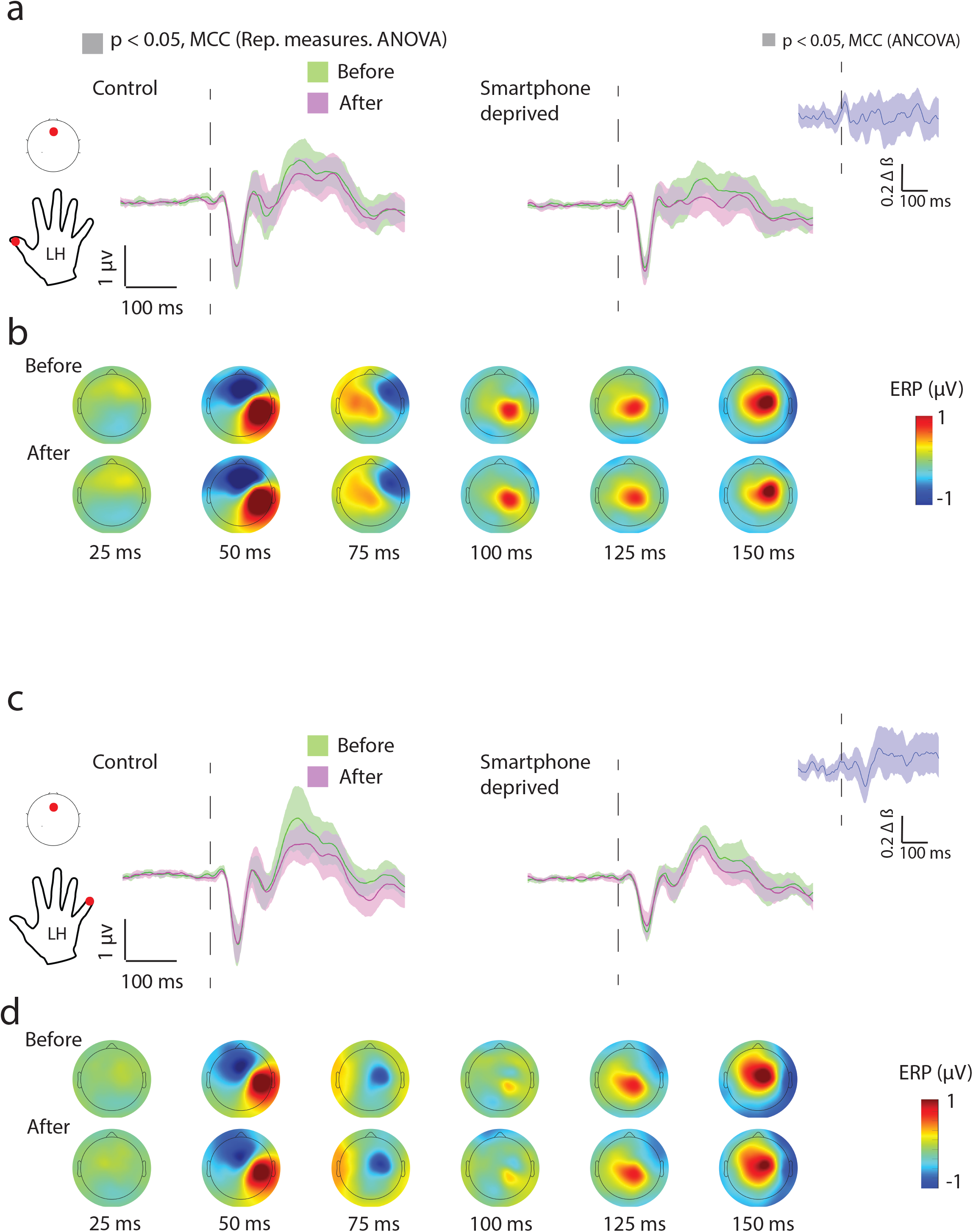
Event-related potentials corresponding to the left hand. The figure layout is the same as in main Figure 2 with the exception that there were no statically significant clusters to report (ANOVA or ANCOVA).

**Supplementary Figure 2.**
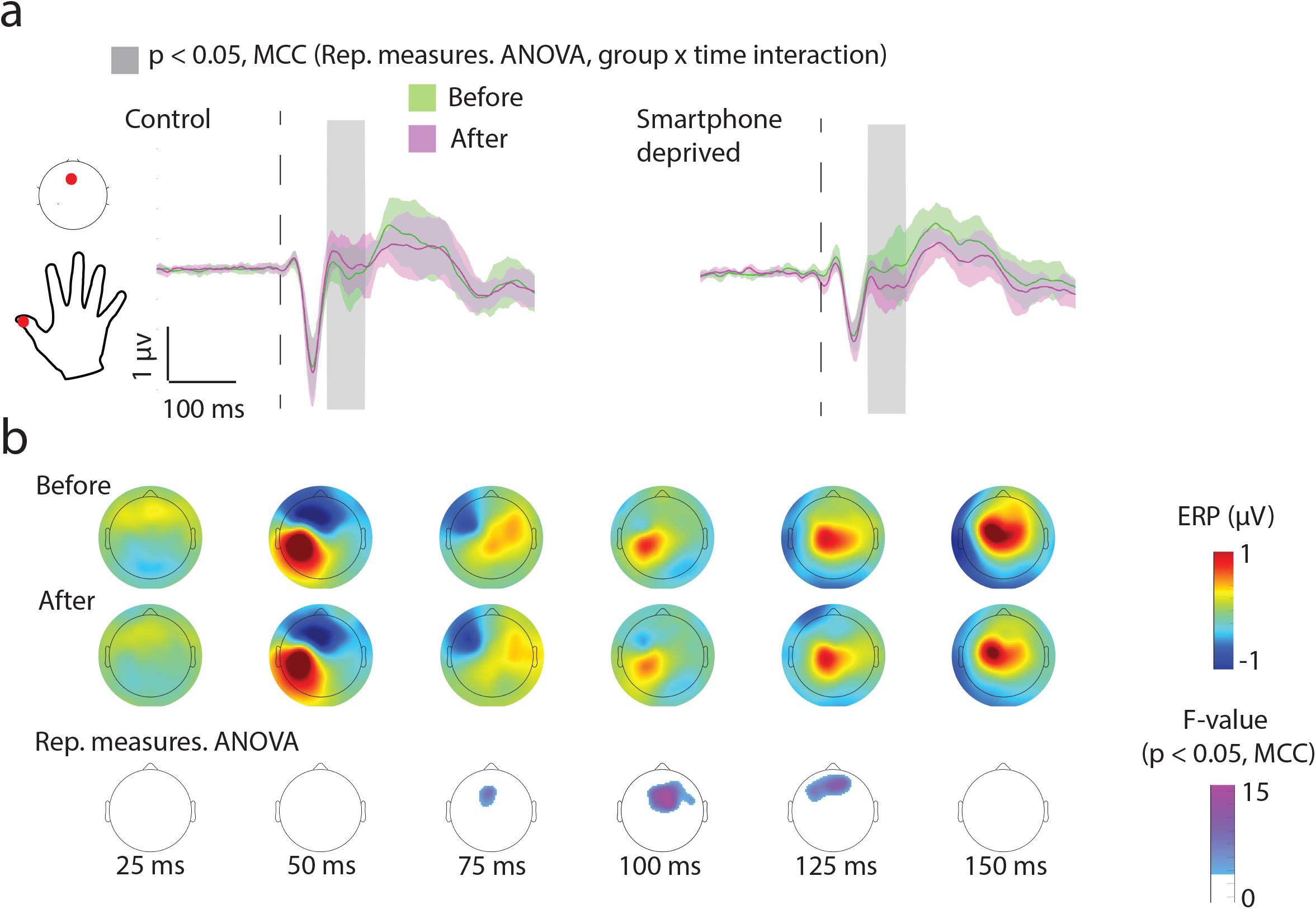
Event-related potentials corresponding to the right hand measured at physical rest. This analysis only included the right thumbed smartphone users determined by using self-reports (hand preference > 1 & thumb preference = 1, see the main Figure. 1). **(a)** Tactile stimulation at the thumb tip results in smaller amplitude EEG signals after smartphone deprivation. Mean (trimmed) and the corresponding 95% confidence intervals are shown from a midline frontal electrode (red dot) with the period of significant group × time interaction clusters (according to repeated measures ANOVA) shaded in grey. **(b)** Scalp plots show mean signals from the smartphone deprivation group and their corresponding interaction *F* values from the ANOVA.

**Supplementary Figure 3.**
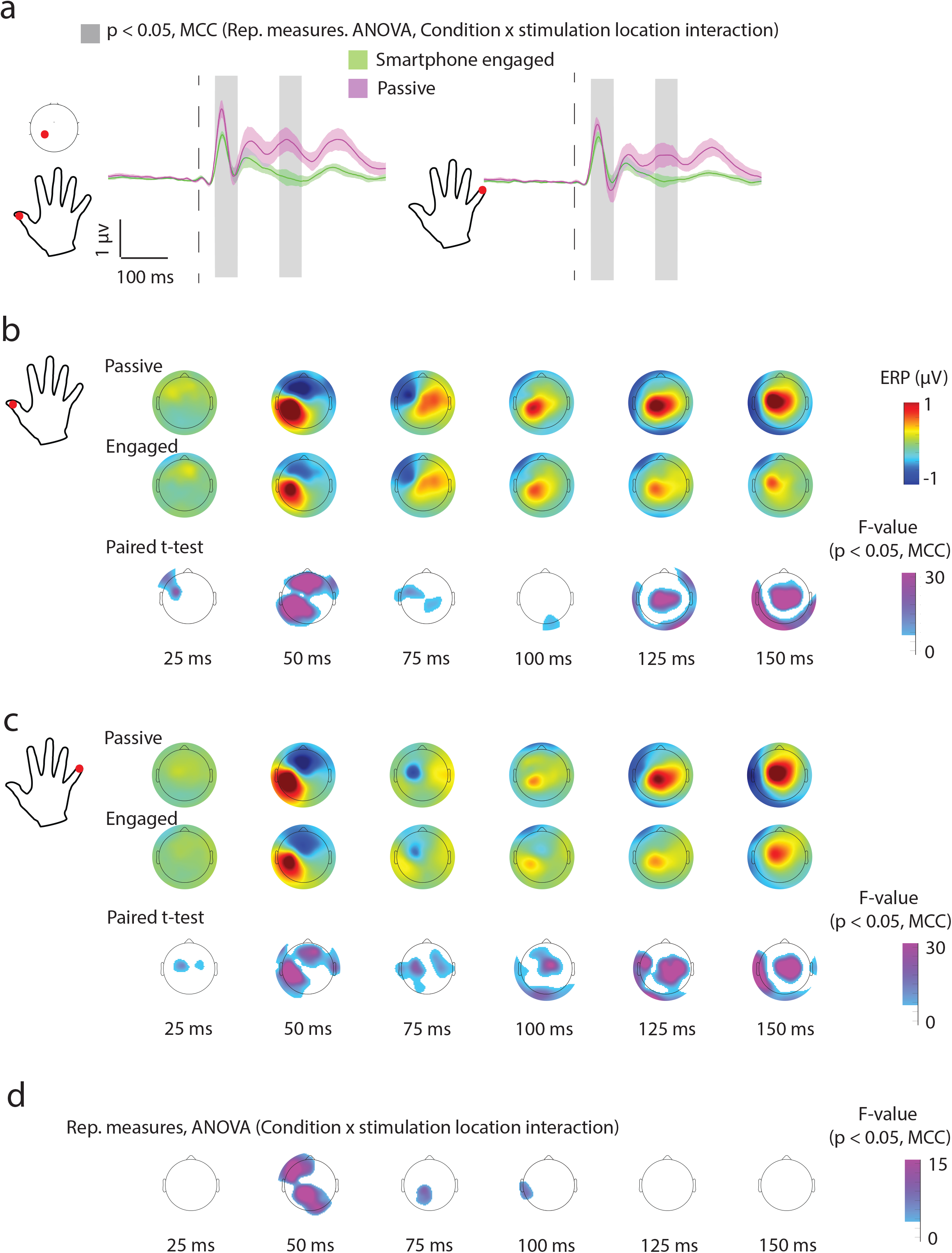
Suppression of tactile sensory signals during smartphone use. **(a)** The tactile event-related potentials from an electrode contralateral to the right thumb and right little finger stimulations. The trimmed means and the corresponding 95% confidence intervals. The shaded area in grey marks the significant repeated measures ANOVA clusters for stimulation location (thumb vs. little finger) × condition (physical rest vs. smartphone engaged) interaction. **(b)** The mean signal amplitudes and the paired *t*-test comparing the different conditions for the thumb. **(c)** The mean signal amplitudes and the paired *t*-test comparing the different conditions for the little finger. **(d)** Repeated Measures ANOVA *F* values showing significant location × condition interaction.

**Supplementary Figure 4.**
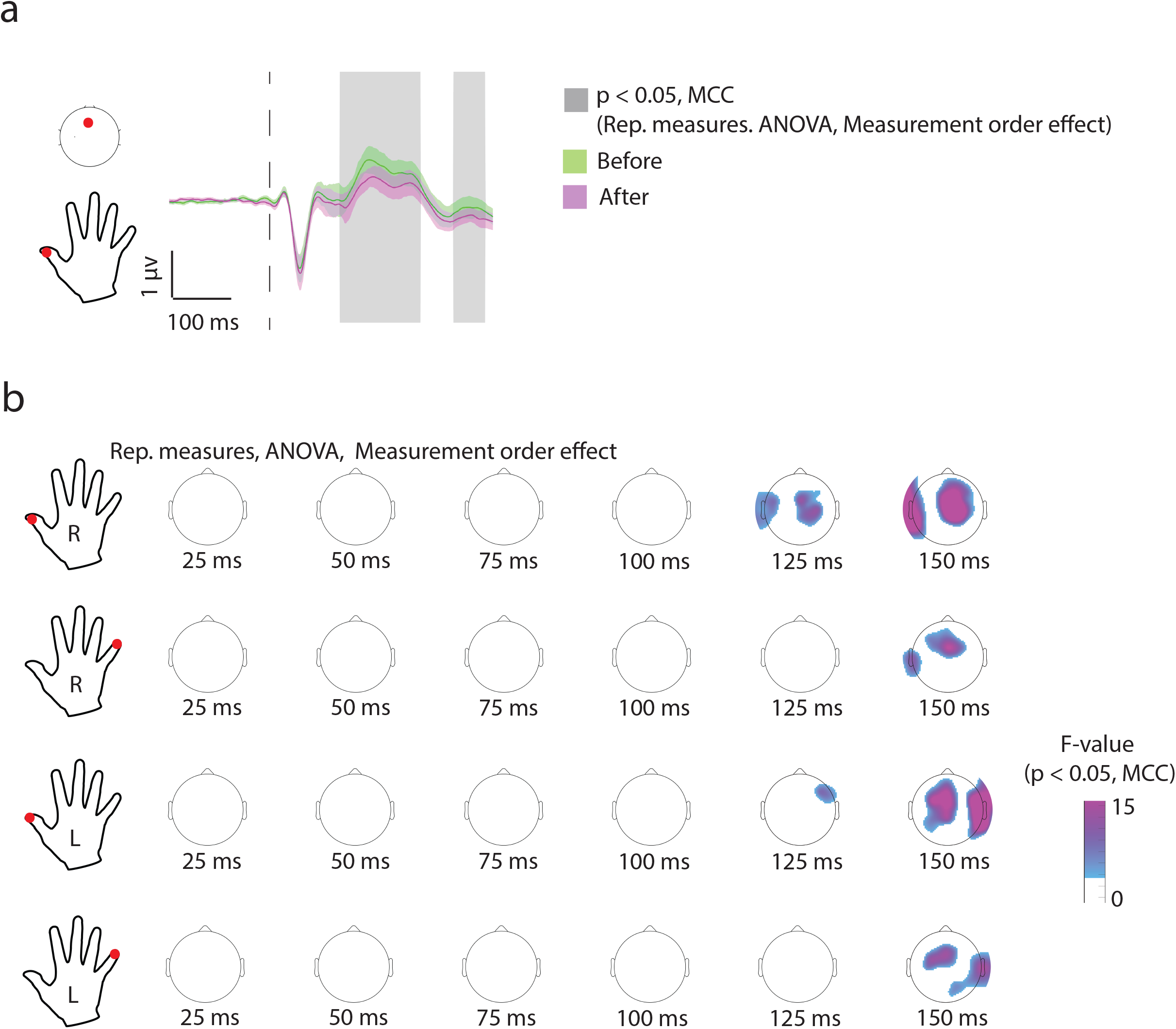
Repetition suppression of EEG signals between sessions separated by several days. **(a)** The mid to late latency signals were suppressed during the second measurement session in comparison to the first. **(b)** This pattern of suppression was visible for all of the stimulated locations at physical rest. No significant clusters were found when this analysis was performed using the tactile stimulations during smartphone use.

## Methods

### Participants

Volunteers were recruited from the university community at Leiden University. Eighty-eight right-handed individuals (self-assessed) were recruited for a study addressing a broad number of questions surrounding smartphone usage (44 females, 16–45 years of age, mean age 23)^11,38^. Only users with a non-shared Android smartphone were included. Moreover, candidates with known neurological or psychiatric conditions (self-assessed) were excluded during recruitment. In 5 participants the background app (see below) failed to operate and 8 participants failed to complete the two measurement sets necessary for this study (yielding a working sample of 75 participants). The participants were rewarded with cash or university credit upon successful completion of the two (before & after) measurement sessions. All participants provided written informed consent and all of the procedures were approved by the Ethical Committee at the Institute of Psychology at Leiden University.

### App-based monitoring of smartphone interactions

The recruited participants were instructed to install an app (TapCounter, QuantActions GmbH, Lausanne) that logged the touchscreen interactions by operating in the background. The Appregistered the timestamp of any event on the touchscreen. These events were sent to the local computers at Leiden University via a cloud-based service operated by QuantActions. The data was subsequently parsed using MATLAB (Mathworks, Natick). Daily tapping speed was derived by using the fastest (first quartile) inter-touch intervals pooled across usage sessions (defined as the period between the screen on & off). A minimum of within-session interaction count of > 10 had to be met for the intervals to contribute to the daily estimate.

### Smartphone deprivation

Before the first measurement, all participants had the background app installed for ~21 days. The median number of daily interactions in this period was used to formulate the instruction for the deprivation. In the randomly allocated deprivation group participants were instructed to use their smartphones less than 1/10^th^ of the typical usage. The app displayed the daily interaction counts and participants were instructed to check the count at the end of each day. The deprivation was instructed to remain until the second measurement session after which the participants could freely use their smartphones. The participants were encouraged to keep the background app running for an additional week after the second measure. Participants were rewarded at the time of the second laboratory measurement session.

### Smartphone usage questionnaire and estimating sleep through the deprivation

To assess which fingers were typically used on the smartphone participants were instructed to rank a set of 18 images from 1 to 18 reflecting their typical usage (https://github.com/codelableidenvelux/Surveys). A rank of 1 indicated the most frequently used posture and a rank of 18 indicated the least used. The images contained 18 different postures including the thumbs, index, ring, and little fingers of either hand. These ranks were re-scaled between 0 and 1 by dividing the original ranks by 18 and then inverted so the best rank corresponds to 1. To quantify hand preference on the smartphone the maximum rank given to a finger posture involving the right hand was divided by the maximum rank for the left hand. To quantify finger preferences the maximum rescaled rank for each finger was retained. Nineteen of the 75 individuals did not provide legible entries or did not complete the survey to use the ranking method described above.

We established sleep durations using actigraphy based on wrist-worn wearables (GENEACTIV, Activinsights, Cambridgeshire) as described before^38^. Briefly, all participants were continuously measured through the study period but 5 participants were eliminated from the sleep analysis as they could not comply with the wearable or the recording malfunctioned. The Cole– Kripke algorithm derived the sleep durations from the recorded accelerations based on a pre-existing implementation on MATLAB^38–40^.

### Inter-trial motor variability in tactile reaction times

Motor variability was extracted from a tactile reaction time task as described before^18^. In brief, participants responded to a tactile stimulation at the left or right thumb tip with a rapid microswitch press. The stimulation consisted of a computer-controlled solenoid tactile stimulator (Tactor, Dancer Design, Merseyside) activated for a 10 ms square wave. The trials were separated by uniformly distributed intervals from 1.2 s to 2.2 s. The left thumb and right thumb were probed in separate blocks consisting of 400 trials per hand. Participants were given a drink break after the first block. The block order was randomized across participants and measurement sessions. The tactile stimulation event, the microswitch press, and release events were digitized using the EEG setup (see below). The time taken to press and then release the microswitch (movement time) was recorded at each trial and the resulting distribution of movement time values was fitted with an ex-Gaussian distribution to determine the variance of the Gaussian part (σ)^41^. One participant who did not complete the measurement due to technical failure was eliminated from the analysis.

### Tactile stimulation at physical rest

Participants were seated on a reclining chair with the arms rested on a flat cushion. The arms were kept at a distance from each other and prevented from crossing over. The participants were instructed to view a distant screen displaying David Attenborough’s Africa series. The participants also wore earphones delivering white noise in addition to the narrators’ commentary. The solenoid tactile stimulators (see above) was attached to the thumb and little finger of the left and right hands. The solenoid activations occurred in a random order across the fingertips with the following features: subsequent touches did not repeat at the same location, the solenoid was active for 10 ms and the inter-touch interval was uniformly distributed between 0.75 ms to 1 s. The participants were stimulated with 1000 solenoid activations for each location. The activation events were digitized using the EEG equipment (see below).

### Tactile stimulation during smartphone use

The thumb and little fingertip of the right hand were stimulated while the participants interacted with their smartphone. Participants were instructed to use their smartphone while operating the two most used (based on their usage history) social and non-social Apps for 15 min per App at a stretch. The participants were instructed to use their right thumb and this use was confirmed by the experimenter using a display connected to a bend-sensitive resistor attached to the thumb. The solenoid stimulator attached to the thumb tip was further wrapped with an aluminum conductor so that the solenoid-covered thumb tip could be used to activate the capacitive touchscreen sensor. The solenoid activations occurred in random order with the following features: the solenoid was active for 10 ms and the inter-touch interval was uniformly distributed between 0.75 ms to 1 s. The participants were stimulated with 1000 solenoid activations per location. The solenoid activation events were digitized using the EEG equipment (see below). Through this period participants were explicitly instructed to not engage with videos or audio functions to ensure a continued touch screen engagement.

### EEG data acquisition and processing

EEG data were gathered using a 64-channel DC amplifier BrainAmp (Brain Products GmbH, Gilching). The EEG caps consisted of passive electrodes with 62 equidistant scalp electrodes and two ocular electrodes (EasyCap, Herrsching). The signals were recorded and digitized at 1 kHz. All of the EEG data processing was performed offline using EEGLAB running on MATLAB^42^.

The data was bandpass filtered between 0.1 and 70 Hz, and data channels reaching > 10 kΩ during the recording period were eliminated. Next, we conducted an independent component analysis for artifact rejection. Eye blinks were rejected based on these components and their correlation to the ocular electrodes by using *icablinkmetrics* implemented in MATLAB^43^. Rejected channels were interpolated and the data were additionally low pass filtered at 30 Hz. Subsequently, an average reference was imposed based on the scalp electrodes. In the final stages of data processing after time-locking, the signals to the tactile stimulation events the trials were baseline corrected according to a baseline window of −200 ms to −1 ms from the stimulation. Time-locked trials exceeding a signal amplitude of 80 μV were eliminated before statistical analysis.

### Statistical analysis of behavioral and EEG data

For the behavioral data derived from the smartphone and for testing the impact of the deprivation Repeated measures, ANOVA was used (implemented in MATLAB). The impact of the deprivation was based on the group × time interaction statistics of the ANOVA model.

The statistical analysis of EEG data was conducted using the linear modeling toolbox LIMO EEG^44^. First, from each measurement session, the event-related potential was estimated using the trimmed mean (20% trimmed). Next, sessions with noisy baselines were eliminated using a 0.16 μV mean signal amplitude threshold applied on the baseline period (−200 ms to −10 ms, based on the 75^th^ percentile values obtained from the thumb stimulation at physical rest) over the electrodes with absolute Θ coordinates < 90. Repeated measures ANOVA was conducted using LIMO EEG at each electrode and time points (absolute Θ coordinates < 135, yielding 60 electrodes on the scalp and from −200 ms to 400 ms from the stimulation). Trimmed population means were used towards ANOVA and the statistics were corrected for multiple comparisons using 1000 bootstraps under H0 followed by Spatio-temporal clustering (α = 0.05). A parallel set of analysis was conducted using gain scores ANCOVA on LIMO EEG with the added variables of the time elapsed between the two measurements, the median tapping speed across the days before the first measurement session, the median number of interactions across the days before the first measurement, and the participant’s gender. Reference of gain score sessions with noisy baselines was eliminated using a 0.16 μV mean signal amplitude threshold applied on the baseline period (−200 ms to −10 ms) over the electrodes with absolute Θ coordinates < 90. One participant with a gap of > 20 days (55 days, 99^th^ percentile) between the two measurement sessions was eliminated along with a subject with unexplained large amplitude signal spikes over the somatosensory areas. The statistics were corrected for multiple comparisons as mentioned above. The 95% confidence intervals used to represent the ANOCA beta values were based on 1000 bootstraps under H1. To compare the signals between the physical rest versus the signals obtained during smartphone use, paired t-tests at each electrode and time point were conducted using LIMO EEG and subsequently corrected for multiple comparisons as mentioned above.

### Date and Code Availability

All of the statistical models, the corresponding data and the functions used to generate the figures will be deposited at dataverse.nl upon peer-reviewed publication.

## References

1. Karl, A., Birbaumer, N., Lutzenberger, W., Cohen, L. G. & Flor, H. Reorganization of Motor and Somatosensory Cortex in Upper Extremity Amputees with Phantom Limb Pain. J. Neurosci. 21, 3609–3618 (2001).

2. Makin, T. R. et al. Deprivation-related and use-dependent plasticity go hand in hand. eLife 2, e01273 (2013).

3. Donoghue, J. P. Plasticity of adult sensorimotor representations. Curr. Opin. Neurobiol. 5, 749–754 (1995).

4. Elbert, T. & Rockstroh, B. Reorganization of Human Cerebral Cortex: The Range of Changes Following Use and Injury. The Neuroscientist 10, 129–141 (2004).

5. Langer, N., Hänggi, J., Müller, N. A., Simmen, H. P. & Jäncke, L. Effects of limb immobilization on brain plasticity. Neurology 78, 182–188 (2012).

6. Taub, E. The Behavior-Analytic Origins of Constraint-Induced Movement Therapy: An Example of Behavioral Neurorehabilitation. Behav. Anal. 35, 155–178 (2012).

7. Huber, R. et al. Arm immobilization causes cortical plastic changes and locally decreases sleep slow wave activity. Nat. Neurosci. 9, 1169–1176 (2006).

8. Crews, R. T. & Kamen, G. Motor-Evoked Potentials Following Imagery and Limb Disuse. Int. J. Neurosci. 116, 639–651 (2006).

9. Moisello, C. et al. Short-Term Limb Immobilization Affects Motor Performance. J. Mot. Behav. 40, 165–176 (2008).

10. Newbold, D. J. et al. Plasticity and Spontaneous Activity Pulses in Disused Human Brain Circuits. Neuron 107, 580–589.e6 (2020).

11. Huber, R. & Ghosh, A. Large cognitive fluctuations surrounding sleep in daily living. iScience 24, (2021).

12. Althoff, T., Horvitz, E., White, R. W. & Zeitzer, J. Harnessing the Web for Population-Scale Physiological Sensing: A Case Study of Sleep and Performance. in Proceedings of the 26th International Conference on World Wide Web 113–122 (International World Wide Web Conferences Steering Committee, 2017). doi:10.1145/3038912.3052637.

13. Min, J.-K. et al. Toss ‘N’ Turn: Smartphone As Sleep and Sleep Quality Detector. in Proceedings of the SIGCHI Conference on Human Factors in Computing Systems 477–486 (ACM, 2014). doi:10.1145/2556288.2557220.

14. Dayan, E. & Cohen, L. G. Neuroplasticity Subserving Motor Skill Learning. Neuron 72, 443–454 (2011).

15. Elbert, T., Pantev, C., Wienbruch, C., Rockstroh, B. & Taub, E. Increased Cortical Representation of the Fingers of the Left Hand in String Players. Science 270, 305–307 (1995).

16. Scholz, J., Klein, M. C., Behrens, T. E. J. & Johansen-Berg, H. Training induces changes in white-matter architecture. Nat. Neurosci. 12, 1370–1371 (2009).

17. Gindrat, A.-D., Chytiris, M., Balerna, M., Rouiller, E. M. & Ghosh, A. Use-Dependent Cortical Processing from Fingertips in Touchscreen Phone Users. Curr. Biol. 25, 109–116 (2015).

18. Balerna, M. & Ghosh, A. The details of past actions on a smartphone touchscreen are reflected by intrinsic sensorimotor dynamics. Npj Digit. Med. 1, 4 (2018).

19. Yoon, J.-S., Ericsson, K. A. & Donatelli, D. Effects of 30 Years of Disuse on Exceptional Memory Performance. Cogn. Sci. 42, 884–903 (2018).

20. Jr, W. A., Jr, W. B., Stanush, P. L. & McNelly, T. L. Factors That Influence Skill Decay and Retention: A Quantitative Review and Analysis. Hum. Perform. 11, 57–101 (1998).

21. Ford, J. K., Baldwin, T. T. & Prasad, J. Transfer of Training: The Known and the Unknown. Annu. Rev. Organ. Psychol. Organ. Behav. 5, 201–225 (2018).

22. Demet, K., Martinet, N., Guillemin, F., Paysant, J. & André, J.-M. Health related quality of life and related factors in 539 persons with amputation of upper and lower limb. Disabil. Rehabil. 25, 480–486 (2003).

23. Grill-Spector, K., Henson, R. & Martin, A. Repetition and the brain: neural models of stimulus-specific effects. Trends Cogn. Sci. 10, 14–23 (2006).

24. Mooney, R. Neural mechanisms for learned birdsong. Learn. Mem. 16, 655–669 (2009).

25. Proteau, L., Marteniuk, R. G. & Lévesque, L. A Sensorimotor Basis for Motor Learning: Evidence Indicating Specificity of Practice. Q. J. Exp. Psychol. Sect. A 44, 557–575 (1992).

26. Peters, M. Why the preferred hand taps more quickly than the non-preferred hand: Three experiments on handedness. Can. J. Psychol. Can. Psychol. 34, 62–71 (1980).

27. Todor, J. I. & Kyprie, P. M. Hand Differences in the Rate and Variability of Rapid Tapping. J. Mot. Behav. 12, 57–62 (1980).

28. Hammond, G. Correlates of human handedness in primary motor cortex: a review and hypothesis. Neurosci. Biobehav. Rev. 26, 285–292 (2002).

29. Jäncke, L., Schlaug, G. & Steinmetz, H. Hand Skill Asymmetry in Professional Musicians. Brain Cogn. 34, 424–432 (1997).

30. Hari, R. et al. Functional Organization of the Human First and Second Somatosensory Cortices: a Neuromagnetic Study. Eur. J. Neurosci. 5, 724–734 (1993).

31. Pleger, B. et al. Functional Imaging of Perceptual Learning in Human Primary and Secondary Somatosensory Cortex. Neuron 40, 643–653 (2003).

32. Chao, L. L., Weisberg, J. & Martin, A. Experience-dependent Modulation of Category-related Cortical Activity. Cereb. Cortex 12, 545–551 (2002).

33. van Turennout, M., Bielamowicz, L. & Martin, A. Modulation of Neural Activity during Object Naming: Effects of Time and Practice. Cereb. Cortex 13, 381–391 (2003).

34. Voss, M., Ingram, J. N., Haggard, P. & Wolpert, D. M. Sensorimotor attenuation by central motor command signals in the absence of movement. Nat. Neurosci. 9, 26–27 (2006).

35. Chapman, C. E., Bushnell, M. C., Miron, D., Duncan, G. H. & Lund, J. P. Sensory perception during movement in man. Exp. Brain Res. 68, 516–524 (1987).

36. Glazewski, S. & Fox, K. Time course of experience-dependent synaptic potentiation and depression in barrel cortex of adolescent rats. J. Neurophysiol. 75, 1714–1729 (1996).

37. Pfister, J.-P. & Ghosh, A. Generalized priority-based model for smartphone screen touches. Phys. Rev. E 102, 012307 (2020).

38. Borger, J. N., Huber, R. & Ghosh, A. Capturing sleep–wake cycles by using day-to-day smartphone touchscreen interactions. Npj Digit. Med. 2, 1–8 (2019).

39. Cole, R. J., Kripke, D. F., Gruen, W., Mullaney, D. J. & Gillin, J. C. Automatic sleep/wake identification from wrist activity. Sleep 15, 461–469 (1992).

40. Sano, A. & Picard, R. W. Toward a taxonomy of autonomic sleep patterns with electrodermal activity. Conf. Proc. Annu. Int. Conf. IEEE Eng. Med. Biol. Soc. IEEE Eng. Med. Biol. Soc. Annu. Conf. 2011, 777–780 (2011).

41. Lacouture, Y. & Cousineau, D. How to use MATLAB to fit the ex-Gaussian and other probability functions to a distribution of response times. Tutor. Quant. Methods Psychol. 4, 35–45 (2008).

42. Delorme, A. & Makeig, S. EEGLAB: an open source toolbox for analysis of single-trial EEG dynamics including independent component analysis. J. Neurosci. Methods 134, 9–21 (2004).

43. Pontifex, M. B., Miskovic, V. & Laszlo, S. Evaluating the efficacy of fully automated approaches for the selection of eyeblink ICA components. Psychophysiology 54, 780–791 (2017).

44. Pernet, C. R., Chauveau, N., Gaspar, C. & Rousselet, G. A. LIMO EEG: A Toolbox for Hierarchical LInear MOdeling of ElectroEncephaloGraphic Data. Computational Intelligence and Neuroscience https://www.hindawi.com/journals/cin/2011/831409/ (2011) doi:10.1155/2011/831409.

